# The transitivity of the Hardy-Weinberg law

**DOI:** 10.1101/2021.09.09.459657

**Authors:** Jan Graffelman, Bruce S. Weir

## Abstract

The reduction of multi-allelic polymorphisms to variants with fewer alleles, two in the limit, is addressed. The Hardy-Weinberg law is shown to be transitive in the sense that a multi-allelic polymorphism that is in equilibrium will retain its equilibrium status if any allele together with its corresponding genotypes is deleted from the population. Similarly, the transitivity principle also applies if alleles are joined, which leads to the summation of allele frequencies and their corresponding genotype frequencies. These basic polymorphism properties are intuitive, but they have apparently not been formalized or investigated. This article provides a straightforward proof of the transitivity principle, and its usefulness in practical genetic data analysis with multi-allelic markers is explored. In general, results of statistical tests for Hardy-Weinberg equilibrium obtained with polymorphisms that are reduced by deletion or joining of alleles are seen to be consistent with the formulated transitivity principle. We also show how the transitivity principle allows one to identify equilibrium-offending alleles, and how it can provide clues to genotyping problems and evolutionary changes. For microsatellites, which are widely used in forensics, the transitivity principle implies one expects similar results for statistical tests that use length-based and sequence-based alleles. High-quality autosomal microsatellite databases of the US National Institute of Standards and Technology are used to illustrate the use of the transitivity principle in testing both length-based and sequence-based microsatellites for Hardy-Weinberg proportions. Test results for Hardy-Weinberg proportions for the two types of microsatellites are seen to be largely consistent and can detect allele imbalance.

## 1 Introduction

The Hardy-Weinberg law is a cornerstone principle of modern genetics, and marked the foundation of population genetics (Crow, 1988). For an autosomal diploid variant, the principle establishes that genotype frequencies attain a stable composition in one generation of time; remaining, in the absence of disturbing forces, unaltered afterwards. For bi-allelic variants this implies the genotype frequencies will have relative frequencies (*AA* = *p*^2^, *AB* = 2*pq, BB* = *q*^2^), where *p* and *q* are the allele frequencies of A and B respectively with *p* + *q* = 1. The Hardy-Weinberg principle becomes more complicated if one considers, for example, X chromosomal variants (Crow and Kimura, 1970), systems with multiple alleles (Hernández and Weir, 1989; Guo and Thompson, 1992; Huber et al., 2006; Aoki, 2003), systems with null alleles (Carlson et al., 2006; McCarroll et al., 2006), copy number variation (Lee et al., 2008; Recke et al., 2015) or polyploid species (Sun et al., 2021; Meirmans et al., 2018). The statistical methodology needed to address all these complications often lags behind, as exemplified by the fact that adequate statistical procedures for testing X chromosomal variants have only been recently developed (Graffelman and Weir, 2016, 2018). In forensics the Hardy-Weinberg law plays a crucial role in probability calculations (Evett and Weir, 1998) and in the analysis of microsatellite data. A basic model forensic genetic model, the Balding-Nichols model, assumes Hardy-Weinberg proportions (HWP) in subpopulations (Balding and Nichols, 1995). In forensic studies, statistical tests for HWP are routinely applied to autosomal microsatellites (STRs (Guevara et al., 2021; Hill et al., 2013)), indels (He et al., 2019), sequence-based STRs (Silva et al., 2020), SNP panels (Simayijiang et al., 2019; Wu et al., 2019) and microhaplotypes (MHs; (Oldoni et al., 2020)) as part of quality control procedures. The analysis of STR data is often complicated by the existence of genotyping error and individuals that stem from different ethnicities or ancestries. The Hardy-Weinberg law is transitive in the sense that it carries over to reduced polymorphisms that can be generated from STRs by elimination or joining of alleles. For STRs, next generation sequencing has revealed additional sequence diversity (Gettings et al., 2018; Silva et al., 2020), thereby increasing the number of STR alleles. Sequence-based (SB) STRs can always be reduced to length-based (LB) STRs. If the equilibrium assumption holds true, one expects inferences made on HWP with SB and LB STRs to be consistent. The main point of this article is that transitivity can be exploited to analyse STR data in more detail. The structure of this article is as follows. In Section 2 we state the transitivity of the Hardy-Weinberg law, and provide a straightforward proof. In Section 3 we explore and use the transitivity principle in the analysis of a reference STR database. A discussion completes the article.

## 2 Theory

The Hardy-Weinberg law states in essence that genotype frequencies are the product of allele frequencies. Let **p** = (*p*_1_, *p*_2_, …, *p*_*k*_)*′* be the column vector of allele frequencies for a genetic variant with *k* alleles. Let **G** be a *k × k* matrix with genotype frequencies, rows representing male alleles and columns representing female alleles. Then the HW law can be concisely expressed as **G** = **pp***′*. In principle, this formulation distinguishes two subtypes of each heterozygote according to the provenance of the maternal and paternal alleles. In general, such distinction is not needed, and both **G** and **pp***′* can always be folded around the diagonal towards a lower triangular matrix with entries 2*p*_*i*_*p*_*j*_ below the diagonal and entries 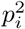 on the diagonal. We here maintain the distinction between the two heterozygote subtypes for mere mathematical convenience, such that **G** = **pp***′* is a sufficient condition for HWP to hold. Assume a population to be in HWP. Intuitively, one might expect that if one allele and its corresponding genotypes are deleted from this population, the law will continue to hold for the reduced array of genotypes. This is indeed true, and a formal demonstration of the property is given below. We call the law *transitive under elimination of alleles* because the equilibrium property is “carried over” to the reduced population with *k −* 1 alleles. Obviously, the process of deleting an allele and its genotypes can be repeated, and this implies that the genotypes of *any* subsystem of *i < k* alleles of an equilibrium system will always be in HW proportions. This transitivity under allele deletion is the theoretical underpinning for the default recoding of multi-allelic variants as bi-allelic in the widely used PLINK software (Purcell et al., 2007).

In genome-wide association studies, variants with multiple alleles are often recoded as bi-allelic variants, with the main goal of enabling the analyst to use available statistical methodology for the analysis of biallelic variants for all variants available in the database. The recoding can however, be carried out in various ways. If all alleles beyond the two most common ones are recoded as missing values, then the foregoing implies that such variants, if in equilibrium, will retain this status. However, elimination of alleles implies a loss of data, leading to smaller sample sizes and less power. The question arises to what will happen if alleles are grouped somehow. To create bi-allelic variants, a straightforward approach is to retain the major allele, and group all remaining alleles as non-major. It is shown below that the law is also *transitive under joining of alleles*.

### Elimination of alleles

We first consider reduction by elimination of alleles. Let **e**_*i*_ be a set of *i* = 1, …, *m* column vectors, where each **e**_*i*_ has one single 1, and all remaining elements equal to zero. We define the *m × k* selector matrix **S**, *m < k*, with a single 1 in each row and at most one 1 in each column, given by

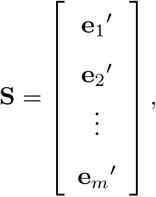

It holds that **SS***′* = **I**_*m*_, and the operation **Sp** removes *k − m* alleles. The vector of reduced and normalized allele frequencies **p**_*r*_ is given by

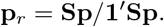

Let **G**_*r*_ represent the reduced matrix of genotype frequencies, where all genotype frequencies that are carriers of the removed allele have been eliminated, and the remaining entries have been renormalized to sum to one. That is

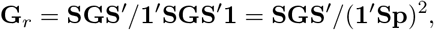

since **1**′**SGS**′**1** = **1**′**Spp**′**S**′**1** = (**1**′**Sp**)^2^. As an example, the reduction of a tri-allelic (A, B, C) to a bi-allelic (A, C) is described by

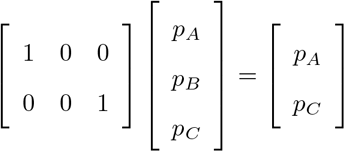

and

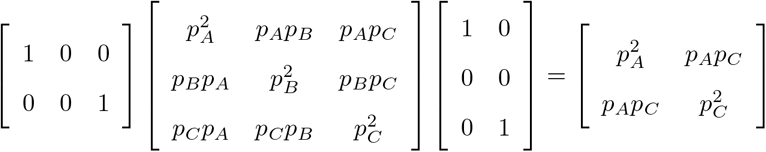

Demonstrating transitivity amounts to showing that the reduced genotype frequencies still satisfy

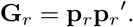

We have

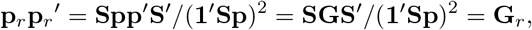

and transitivity is thus established.

### Joining of alleles

We also consider the reduction of the polymorphism by joining alleles, summing the corresponding allele and genotype frequencies. Joining alleles into a single, more frequent allele can be done by a selecting and summing operation on matrix **G**. E.g. if alleles A and B are joined this can be seen as a relabelling of all B alleles as A alleles, such that BB homozygotes and AB heterozygotes become AA homozygotes, and all BC heterozygotes become AC heterozygotes. We define **S** as the *m × k* selector-summing matrix with *m ≤ k*, given by

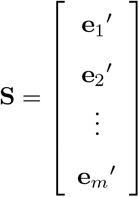

The elements of each vector **e**_*i*_ are either 0 or 1; the matrix has a single 1 in each column and at least a 1 in each row. The vector of reduced and normalized allele frequencies is given by

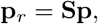

and the *m × m* matrix of reduced and normalized genotype frequencies is given by

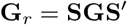

As an example, the reduction of a tri-allelic to a bi-allelic by joining the B and the C alleles is described by:

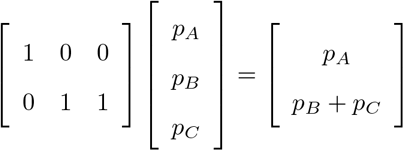

and

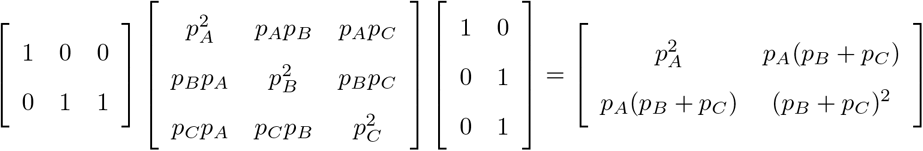

By the same token as before, we have

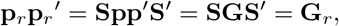

and transitivity is again established. When alleles are joined, the re-normalization to unit-sum allele and genotype frequencies is not required.

## 3 NIST microsatellites

We illustrate the application of the formulated transitivity principle in genetic data analysis with microsatellites. We use a microsatellite database of the NIST website consisting of 1036 individuals of four different self-identified ethnicities genotyped for 29 autosomal length-based (LB) STRs (https://strbase.nist.gov/). This data set has been described by Hill et al. (2013), and posterior corrections are detailed by Steffen et al. (2017). A sequence-based (SB) version of 27 STRs for the same individuals has been described by Gettings et al. (2018); the LB and SB data sets have 23 STRs in common. The NIST data set underwent extensive concordance evaluations and is probably one of most reliable STR databases publicly available. Hill et al. (2013) report on HWP test results, and argue that after using a Bonferroni correction for multiple testing, only two significant deviations remain (D13S317 and F13B when tested overall). Gettings et al. (2018) report no significant deviations from HWP in the SB data after correction for multiple testing.

Here, we present a HWP analysis of the LB and SB STR data exploiting the transitivity principle in various ways. In all cases we test for HWP by using the mid *p*-value, as this has been shown to have a rejection rate that is most close to the nominal level (Graffelman and Moreno, 2013). For bi-allelic variants, we calculate the exact mid *p*-value. For multi-allelic variants, we estimate the mid *p*-value with a permutation test using 17,000 random shuffles of the alleles. This estimates the exact mid *p*-value with a precision of 1% with 99% confidence (Guo and Thompson, 1992; Ziegler and König, 2006). To illustrate transitivity, we first use CSF1PO in the sample of Caucasian ethnicity only. For this STR, seven alleles are observed, and HWP is not rejected (*p*-value = 0.866) in a permutation test that uses the exact probability of the data table according to Levene’s distribution (Levene, 1949) as a test statistic. Table 1 shows mid *p*-values of seven permutation tests for HWP where just one allele is eliminated, each in turn. In all cases, HWP is not rejected for the reduced six-allelic polymorphisms as expected by transitivity. Table 1 also shows the mid *p*-values obtained after a bi-allelic recoding of all alleles (e.g. “8” versus “not-8”), using a standard bi-allelic exact test for HWP (Weir, 1996). This also produces no significant results, as is again expected by transitivity. The results in Table 1 are representative for most STRs of the NIST database, when stratifying for ethnicity (results not shown). When the sequence-based STRs are used, the same results are obtained because the Caucasian sample has no additional sequence variability for this STR.

**Table 1:** Permutation test and bi-allelic test results for HWP of STR CSF1PO of the Caucasian sample. SB- allele: identifier and sequence motif; LB-allele: repeat number; *n*: sample size (number of individuals) after elimination of the allele; *p*: allele frequency; *p*_*mid*_: permutation test mid *p*-value obtained when deleting the allele; AA, AB, BB: genotype counts of the bi-allelified polymorphism (A = minor, retained allele, B = all other alleles); *p*_*mid*_: mid *p*-value of bi-allelic exact test for HWP.

| SB-allele | LB-allele | $n$ | $p$ | $p_{mid}$ | AA | AB | BB | $p_{mid}$ |
| --- | --- | --- | --- | --- | --- | --- | --- | --- |
| – | – | 361 | – | 0.8661 | – | – | – | – |
| 238 [ATCT]8 | 8 | 357 | 0.005 | 0.7570 | 0 | 4 | 357 | 0.5042 |
| 239 [ATCT]9 | 9 | 351 | 0.014 | 0.8463 | 0 | 10 | 351 | 0.5306 |
| 240 [ATCT]10 | 10 | 219 | 0.220 | 0.5810 | 17 | 125 | 219 | 0.9392 |
| 242 [ATCT]11 | 11 | 173 | 0.309 | 0.6892 | 35 | 153 | 173 | 0.8537 |
| 245 [ATCT]12 | 12 | 145 | 0.360 | 0.8520 | 44 | 172 | 145 | 0.5310 |
| 247 [ATCT]13 | 13 | 302 | 0.082 | 0.9198 | 0 | 59 | 302 | 0.1131 |
| 248 [ATCT]14 | 14 | 354 | 0.010 | 0.8336 | 0 | 7 | 354 | 0.5145 |

More interesting is the removal (or amalgamation) of alleles for a variant for which HWP is rejected. A large change in *p*-value, from clearly significant to clearly non-significant after removal can signal which allele(s) provoked the initial rejection of the null. Table 2 shows such test results for the full database with all ethnicities for TH01. For this STR, HWP are rejected in the full database for both the LB and SB data (first row of Table 2). This table shows that the elimination of fractional allele 9.3 renders the test non-significant, suggesting 9.3 is an equilibrium-offending allele, or at least the most offending allele. Figure 1 shows a barplot of the allele frequencies of this STR for the four ethnicities. This reveals the 9.3 allele fluctuates strongly over the ethnicities, ranging from 4% in Asians to 34% in Caucasians. Interestingly, if the 9 and 9.3 alleles are joined, then there is no significant difference in allele frequencies between Asians, Caucasians and Hispanics. An alternative way to identify equilibrium-offending alleles is to calculate the contribution each allele makes to the chi-square statistic of the original data table; in this case the homozygote 9.3 makes the largest contribution. It is well-known that population substructure can drive disequilibrium when there are differences in allele frequencies across the groups. For all length-based and sequence-based STRs in the NIST database, a Fisher exact test for equality of allele frequencies across the four ethnicities is highly significant at a Bonferroni corrected significance level (0.05*/*29 = 0.0017 for LB; 0.05*/*27 = 0.0019 for SB; results not shown). It is thus imperative to account for ethnicity when testing for HWP.

**Table 2:** Test results of a permutation test for HWP of TH01, after removal of a single allele. Allele: removed allele; *p*: allele frequency; *n*: sample size (number of individuals) after elimination of the allele; *p*_*mid*_: mid *p*-value obtained in a permutation test.

| Sequence-based (SB) |  |  |  |  | Length-based (LB) |  |  |  |
| --- | --- | --- | --- | --- | --- | --- | --- | --- |
| ID | Allele | $p$ | $n$ | $p_{mid}$ | Allele | $p$ | $n$ | $p_{mid}$ |
| – | – | – | 1036 | 0.0001 | – | – | 1036 | 0.0003 |
| 379 | [AATG]5 | 0.0019 | 1032 | 0.0001 | 5 | 0.0019 | 1032 | 0.0001 |
| 380 | [AATG]6 | 0.1959 | 669 | 0.0001 | 6 | 0.1959 | 669 | 0.0000 |
| 381 | [AATG]7 | 0.2939 | 524 | 0.0036 | 7 | 0.2949 | 524 | 0.0035 |
| 382 | [AATG]7-rs1051822965 | 0.0010 | 1034 | 0.0004 |  |  |  |  |
| 383 | [AATG]8 | 0.1255 | 791 | 0.0001 | 8 | 0.1255 | 791 | 0.0001 |
| 384 | [AATG]9 | 0.1684 | 729 | 0.0047 | 9 | 0.1689 | 728 | 0.0030 |
| 385 | AGTG [AATG]8 | 0.0005 | 1035 | 0.0001 |  |  |  |  |
| 386 | [AATG]6 ATG [AATG]3 | 0.2056 | 677 | 0.2338 | 9.3 | 0.2056 | 677 | 0.1958 |
| 387 | [AATG]10 | 0.0068 | 1022 | 0.0001 | 10 | 0.0068 | 1022 | 0.0004 |
| 388 | [AATG]11 | 0.0005 | 1035 | 0.0002 | 11 | 0.0005 | 1035 | 0.0002 |

**Figure 1:**
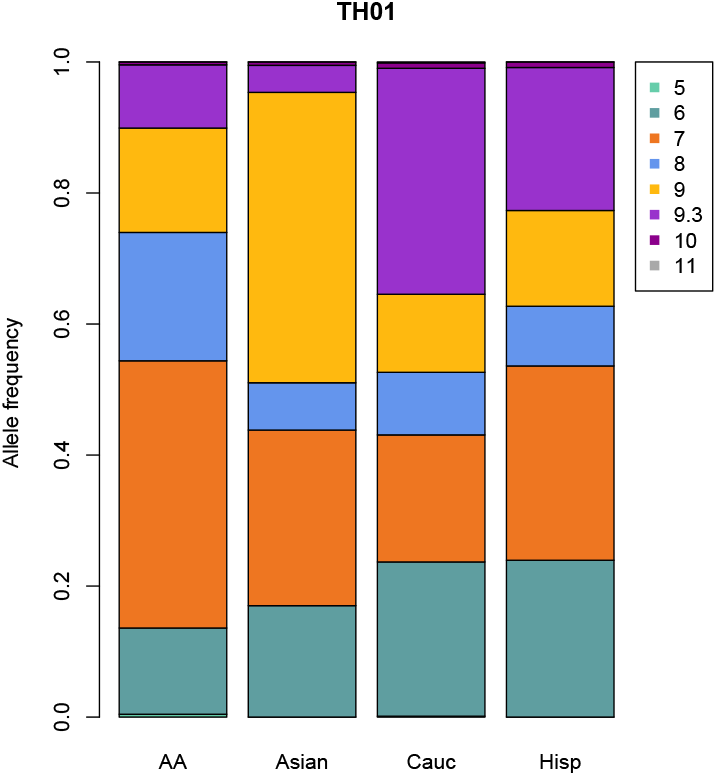
Allele frequencies of TH01 for four ethnicities.

The results obtained for LB and SB alleles in Table 2 are entirely consistent. In both cases the 9.3 allele is identified as equilibrium-offending. The two new alleles generated by using sequences are rare variations on repeats 7 and 9, and their separate elimination does not qualitatively alter the test result.

The testing of all STRs separately for each ethnicity provokes a multiple testing problem. Hill et al. (2013) use a Bonferroni correction taking into account that 29 *×* 4 = 116 tests are being performed, using *α* = 0.05*/*116 = 0.0004. However, the Bonferroni correction is very conservative, and consequently, informative disequilibrium may easily go undetected. The extent of the multiple testing problem can be diminished by using *restricted* permutation test procedures (Manly, 2007) that account for the four-group structure of the data. We applied restricted permutation tests (permuting alleles only within ethnicities) and this allows us to test each STR just once, applying a less restrictive Bonferroni threshold of 0.05/29=0.0017 for the LB data, or 0.05/27=0.0019 for the SB data. At this level we found none of the STRs to be significant, though SE33 is close to the threshold. The test results for all length-based and sequence-based STRs are reported in Table 3. Qualitatively, test results obtained for LB and SB STRs are similar, and among the 23 common ones for which both LB and SB data are available, the two most significant ones are the same, D22S1045 and FGA. When all STRs are considered, SE33 and D22S1045 are the most significant STRs. If we finally test, albeit aggravating the multiple testing problem, each STR for HWP *within* each ethnicity, as is often done, SE33 of the Asian samples singles out as the most significant variant, with permutation mid *p*-value 0.008. Table 4 shows the most significant SB and LB STRs that have a *p*-value below 0.05 and their heterozygosities. D22S1045 and FGA are shared in the SB and LB list. However, SE33 and D4S2408 are the most significant results.

**Table 3:**
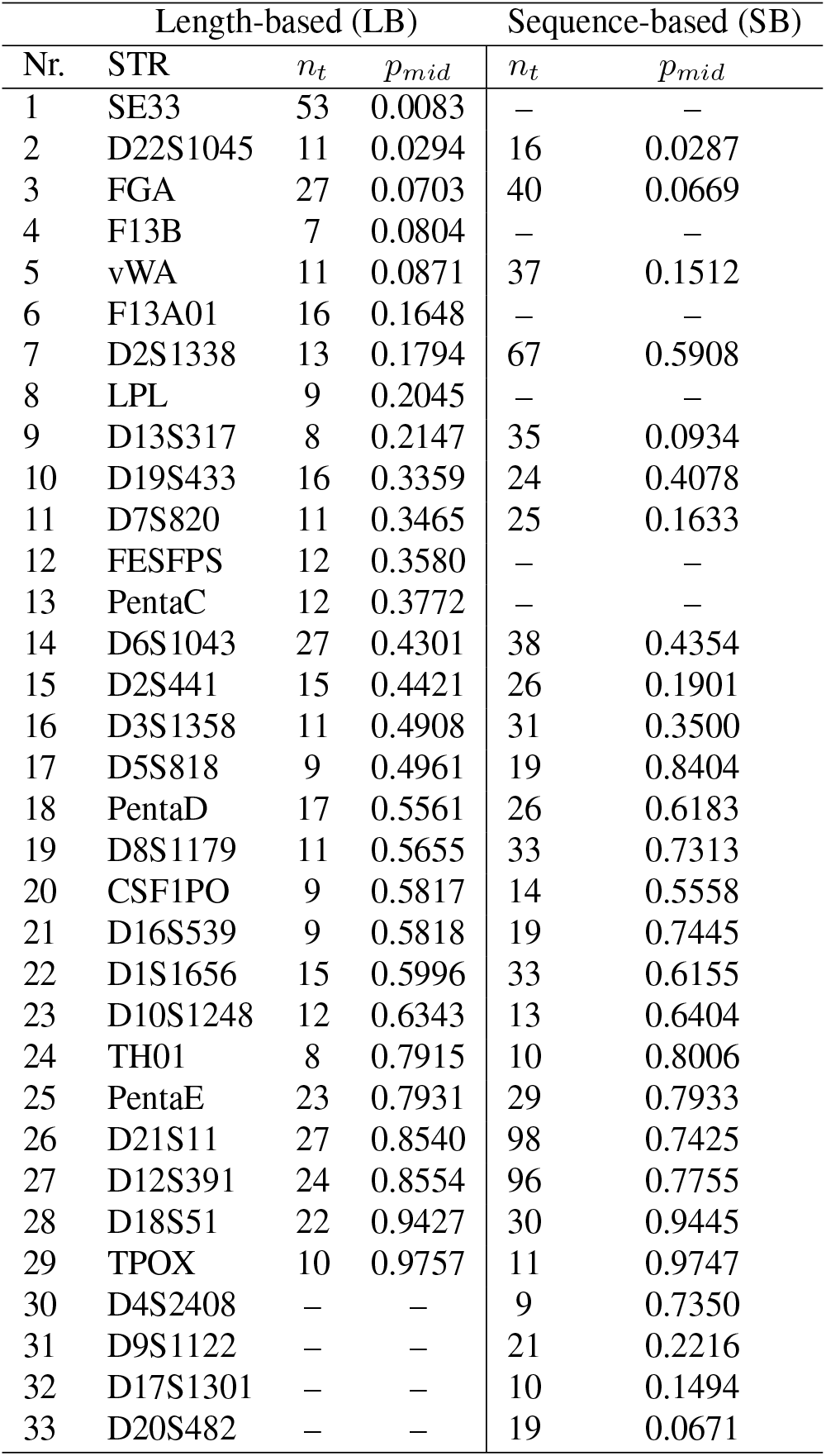
Number of alleles (*n*_*t*_) and mid *p*-values of both length-based and sequence-basted NIST STRs using a restricted permutation test for HWP.

**Table 4:** Results of the most significant permutation tests (*α* = 0.05) for HWP within populations for both LB and SB STRs. STR: microsatellite identifier; *n*: sample size (number of individuals); *p*_*mid*_: mid p-value obtained by permutation; *H*_*o*_: observed heterozygosity; *H*_*e*_: expected heterozygosity.

| Length-based (LB) |  |  |  |  |  | Sequence-based (SB) |  |  |  |  |  |
| --- | --- | --- | --- | --- | --- | --- | --- | --- | --- | --- | --- |
| STR | Population | $n$ | $p_{mid}$ | $H_o$ | $H_e$ | STR | Population | $n$ | $p_{mid}$ | $H_o$ | $H_e$ |
| SE33 | Asian | 97 | 0.0078 | 0.9278 | 0.9392 | D4S2408 | Hispanic | 236 | 0.0021 | 0.6864 | 0.7625 |
| D6S1043 | Cauc | 361 | 0.0217 | 0.7978 | 0.8232 | D5S818 | Hispanic | 236 | 0.0121 | 0.7797 | 0.7874 |
| Penta_C | Cauc | 361 | 0.0229 | 0.7701 | 0.7523 | D22S1045 | Asian | 97 | 0.0272 | 0.7331 | 0.6865 |
| D22S1045 | Asian | 97 | 0.0301 | 0.7010 | 0.7628 | D12S391_1 | Hispanic | 236 | 0.0280 | 0.9576 | 0.9358 |
| D2S441 | Hispanic | 236 | 0.0415 | 0.7542 | 0.7488 | D22S1045 | Hispanic | 236 | 0.0416 | 0.7010 | 0.7628 |
| FGA | Cauc | 361 | 0.0481 | 0.8670 | 0.8594 | FGA | Cauc | 361 | 0.0429 | 0.8670 | 0.8596 |

If we take the results of Tables 3 and 4 together, then the main leads are SE33 and D22S1045; SE33 in the Asian sample, and D22S1045 in both the Asian and Hispanic sample. D221045 has a relatively larger difference between observed and expected heterozygosity; this is also observed for D4S2408 in Hispanics. We follow up these most important leads in an attempt to understand the nature of disequilibrium.

Microsatellite SE33 has many rare fractional alleles. Several rare fractional alleles (18.5, 20.2, 23.2, 28.2) occur in homozygote form, which is very unlikely under the HWP assumption. In fact, if fractional alleles 20.2 and 23.3 alleles are joined with their main repeat, evidence against HWP diminishes (*p* = 0.050). In non-Asian samples there is overall no evidence against HWP, but if SE33 is bi-allelified for its alleles, then 22 and 23.2 have the smallest *p*-values in Hispanics and African Americans again due to the existence of rare allele homozygotes. We tentatively suggest that some of the rare allele homozygotes may in fact be heterozygotes, and that checking for allele imbalance is called for (in particular in the light of the results described for D22S1045 below). We note that Churchill et al. (2016, Fig. 4) report a relatively lower allele coverage ratio (*≈* 0.60) for this STR, and that Just et al. (2017) also reported problems with this locus.

The second most significant STR is D22S1045; there is evidence against HWP for this STR in the Asian sample (*p* = 0.0266), and by deletion of alleles and bi-allelification, repeat 17, a frequent allele, is identified as the most offending allele (see supplementary Table S1; the corresponding table for the SB STR is identical, because no additional sequence variation exists for the Asian sample). The reduced bi-allelic polymorphisms “17” versus “not-17” has a lack of heterozygotes. Investigating the same polymorphism in the other samples shows some evidence against HWP for this STR when it is bi-allelified for allele 16 or 17 (see supplementary Table S2). Most notably, Peng et al. (2020) reported allele imbalance for this STR in a sample from Tibet, with miscalling of heterozygotes as homozygotes, and Novroski et al. (2016) reported heterozygote imbalance for D22S1045 and problems with allele 17 in a US sample.

For D4S2408 we find significant deviations only for the Hispanics; by deleting alleles and bi-allelification, sequence 170 (repeat 9) is identified as equilibrium-offending in Hispanics (results not shown); this pattern is however, not observed in other samples.

## 4 Discussion

We have given a formal proof of the transitivity of the Hardy-Weinberg law, and illustrated its use in genetic data analysis. Only the most simple reductions obtained by elimination of one allele and by bi-allelification have been used. Many additional reductions are possible, such as the reduction to all tri-allelic variants, all four-allelic variants, and so on. In principle, if equilibrium holds true, one expects HWP not to be rejected in most cases, except for chance effects. Reduction to tri-allelics and other multi-allelic forms has not been carried out in order not to further aggravate the multiple testing problem.

Our proof of transitivity is written in plain matrix algebra. However, the operations we performed on genotype and allele frequencies can be rephrased in terms of basic operations known in compositional data analysis (Aitchison, 1986). Genotype and allele frequencies can be considered as compositional data, as both are subject to a unit-sum constraint. The elimination of alleles then corresponds to the creation of a *subcomposition*, the corresponding (re)normalization of allele and genotype frequencies is known as *closure*, and the joining of alleles is known as the *amalgamation* of parts.

For the NIST data, testing each STR within each population using a Bonferroni correction seems too conservative and, as argued above, may leave important disequilibrium undetected. Alternatively, one might use the false discovery rate (Benjamini and Hochberg, 1995) which is less conservative than the Bonferroni correction. We suggest, at any rate, a flexible approach, where even STRs that are strictly speaking not significant but close to the Bonferroni threshold are followed up for inspection of possible causes for disequilibrium. We suggest, as Ye et al.(2020), the source of deviations from HWP to be investigated. Most importantly, we have shown that HWP tests can identify allele imbalance and pinpoint the problematic alleles. The combined inspection of HWP test results and allele coverage ratios seems particularly useful to identify problems. By using a restricted permutation test that permutes alleles only within ethnicities, each STR can be tested for HWP just once, accounting for the fact that allele frequencies can differ over ethnicities. This reduces the multiple testing burden.

Over the last few years, more SB STR data sets have become available for use in forensic studies, and the statistical analysis of the data needs to be adjusted accordingly, as is for instance the case for the study of population substructure (Aalbers et al., 2020). SB STRs have more alleles, and will likely provide additional insights for different topics in population genetics, such as STR mutation and genetic diversity.

Under the assumption of HWP, rare alleles most likely occur in heterozygote form. Significant deviations from HWP easily result if rare alleles occur in homozygote form. We have shown that the grouping of equilibrium-offending alleles can be used to reduce disequilibrium, as this may actually reduce genotyping error if the corresponding alleles that are joined can indeed not be faithfully distinguished. This preserves the assumption of allelic indepence at the small cost of increasing allele frequency and decreasing the numerical strength of matching STR profiles. If there is indication that two alleles can not be distinguished, such as a fractional and a main allele, then it is prudent to combine them. This will preserve the sample size, and strengthen that the HWP requirement of forensic models, the Balding-Nichols model in particular, is met.

## Acknowledgements

This work was partly supported by the Spanish Ministry of Science, Innovation and Universities and the European Regional Development Fund (grant number RTI2018-095518-B-C22 (MCIU/AEI/FEDER)); by the National Institutes of Health (grant number GM075091) and the National Institute of Justice (grant number 2020-DQ-BX-0022). We thank Katherine Gettings and the National Institute of Standards and Technology for making the sequence-based STR data available to us.

## Supplementary tables

**Table S1:**
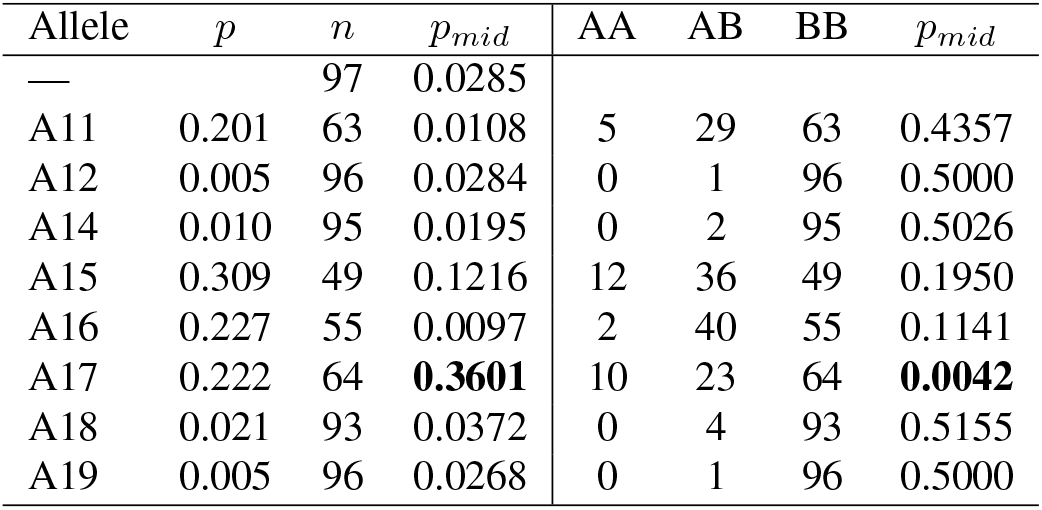
Allele frequency (*p*), sample size (*n* individuals) and mid *p*-value of tests for HWP after deletion of an allele (first four columns) and after bi-allelification (last four columns: genotype counts and mid *p*-value) of the STR D22S1045 of the Asian sample.

**Table S2:** Bi-allelic exact tests results (exact mid *p*-values) for HWP for bi-allelified versions of D22S1045 for four ethnicities.

| Allele | AA |  |  |  | Asian |  |  |  | Caucasian |  |  |  | Hispanic |  |  |  |
| --- | --- | --- | --- | --- | --- | --- | --- | --- | --- | --- | --- | --- | --- | --- | --- | --- |
| | AA | AB | BB | $p_{mid}$ | AA | AB | BB | $p_{mid}$ | AA | AB | BB | $p_{mid}$ | AA | AB | BB | $p_{mid}$ |
| A8 | 0 | 5 | 337 | 0.5073 | — | — | — | — | — | — | — | — | — | — | — | — |
| A10 | 1 | 26 | 315 | 0.2692 | — | — | — | — | — | — | — | — | 0 | 7 | 229 | 0.5221 |
| A11 | 6 | 87 | 249 | 0.7454 | 5 | 29 | 63 | 0.4357 | 7 | 87 | 267 | 0.9135 | 1 | 28 | 207 | 0.8036 |
| A12 | 3 | 31 | 308 | 0.0385 | 0 | 1 | 96 | 0.5000 | 0 | 9 | 352 | 0.5246 | 0 | 6 | 230 | 0.5158 |
| A13 | 0 | 2 | 340 | 0.5007 | — | — | — | — | 0 | 5 | 356 | 0.5069 | 0 | 4 | 232 | 0.5064 |
| A14 | 1 | 51 | 290 | 0.5735 | 0 | 2 | 95 | 0.5026 | 0 | 41 | 320 | 0.4668 | 2 | 9 | 225 | 0.0047 |
| A15 | 23 | 126 | 193 | 0.6167 | 12 | 36 | 49 | 0.1950 | 36 | 160 | 165 | 0.7636 | 39 | 123 | 74 | 0.3198 |
| A16 | 20 | 91 | 231 | **0.0108** | 2 | 40 | 55 | 0.1141 | 44 | 188 | 129 | **0.0515** | 20 | 125 | 91 | **0.0123** |
| A17 | 13 | 117 | 212 | 0.5688 | 10 | 23 | 64 | **0.0042** | 2 | 50 | 309 | 0.8538 | 1 | 41 | 194 | 0.5605 |
| A18 | 0 | 10 | 332 | 0.5323 | 0 | 4 | 93 | 0.5155 | 0 | 4 | 357 | 0.5042 | 0 | 3 | 233 | 0.5032 |
| A19 | 0 | 4 | 338 | 0.5044 | 0 | 1 | 96 | 0.5000 | — | — | — | — | — | — | — | — |

